# PhyD3: a phylogenetic tree viewer with extended phyloXML support for functional genomics data visualization

**DOI:** 10.1101/107276

**Authors:** Łukasz Kreft, Alexander Botzki, Frederik Coppens, Klaas Vandepoele, Michiel Van Bel

## Abstract

**Motivation:** Comparative and evolutionary studies utilise phylogenetic trees to analyse and visualise biological data. Recently, several web-based tools for the display, manipulation, and annotation of phylogenetic trees, such as iTOL and Evolview, have released updates to be compatible with the latest web technologies. While those web tools operate an open server access model with a multitude of registered users, a feature-rich open source solution using current web technologies is not available.

**Results:** Here, we present an extension of the widely used PhyloXML standard with several new options to accommodate functional genomics or annotation datasets for advanced visualization. Furthermore, PhyD3 has been developed as a lightweight tool using the JavaScript library D3.js to achieve a state-of-the-art phylogenetic tree visualisation in the web browser, with support for advanced annotations. The current implementation is open source, easily adaptable and easy to implement in third parties’ web sites.

**Availability:** More information about PhyD3 itself, installation procedures, and implementation links are available at http://phyd3.bits.vib.be and at http://github.com/vibbits/phyd3/.

## 1. Introduction

The construction, visualisation and interpretation of phylogenetic trees are instrumental for biological analysis in multiple fields, including evolutionary biology, genetics, and comparative genomics. Recently published software tools allow for the use of interactive elements and comprehensive analytics in association with these trees by means of the latest web technologies (Letunic and Bork, 2016; He et al., 2016; Smits and Ouverney, 2010; Kuraku et al., 2013). The phyloXML standard (Han and Zmasek, 2009) is widely supported in those tools to visualise the evolutionary trees as well as associated data.

A comparison of the most popular phylogenetic tree viewers reveals that a feature rich open source tool that supports extended charting, is easily adaptable, and can be seamlessly implemented in third party websites, is currently not available (see Supplementary Data). To this end, we extended the phyloXML standard with several new elements in order to accommodate common annotation datasets and developed a lightweight phylogenetic tree viewer using the D3.js JavaScript library (http://d3js.org)

## 2. Methods

The commonly used phyloXML standard has been extended with various elements of the complex type as permitted by the current phyloXML XSD scheme. The newly created elements <taxonomies> and <domains> are used to specify taxonomy and domain colours, descriptions, and links. In order to reference between the <phylogeny> element of the current standard and the new elements, the <code> and <name> subelements are used for the <taxonomy> and the <domain> element, respectively. Colours are represented as HEX values (e.g. 0xRRGGBB). Furthermore, graphs like a pie, a binary, and a multi-bar chart as well as a heat map can be displayed next to the leaf nodes by using the newly defined <graph> elements.

The pie chart and binary chart types can also be associated with internal nodes of the phylogenetic tree. Referencing of the chart values to the nodes is achieved by defining an <id> tag in the <clade> element corresponding to the ‘for’ attribute in <value> elements.

A complete graph specification consists of the following child elements:

1. name: This element contains the graph name that will be displayed in a detailed node information popup.
2. legend: For each value series of your data, a legend field has to be specified. This legend consists of the field name (which will be displayed to the user in the legend and info popup), the colour that will be used to draw this series (for all graph types besides heat map) and an additional symbol shape (e.g. circle) for the binary graph. For the heat map, a gradient specification has to be applied according to the specifications of the ColorBrewer2 project (http://colorbrewer2.org).
3. data: Data values for the graph have to be defined for each clade id for which a graph is displayed. They have to be sorted according to how the series are defined in the respective legend fields. Existing tags or properties from the phylogeny element can be used by referencing them via the ‘tag’ and ‘ref’ attributes of the data element.

## 3. Features

Using the D3.js JavaScript library, PhyD3 is implemented as a flexible and lightweight tool allowing for the display of interactive and complex phylogenetic trees in a web-based environment without security-based limitations and without the need for external plugins. The implementation is fast and responsive to user interaction, with the tree elements’ display parameters being easily changed through user-friendly access controls.

PhyD3 offers a number of key features with respect to tree nodes and graphs visualisation. Basic information (like node and taxonomy names, branch length and support values) can be displayed next to the respective nodes, with detailed information (including link outs to external URLs) being shown in an info box per user request. The display of the information is flexible according to the users’ preferences (e.g. all content, or only internal or external node annotations). To support a clearly annotated data presentation and better readability, leaf nodes can be lined up to match the furthest node and text can be scaled.

Furthermore, nodes can be annotated with structural and numerical information, displayed in form of various graphs. Structural information contains for example the domain or gene structure architecture. PhyD3 provides controls to scale and filter this information to support easy analysis. Numerical annotations are presented using multiple bar charts, pie charts or binary charts. Users can easily change the size and scale of graphs, to match their needs.

Additionally, a number of convenience features were introduced in PhyD3: tree nodes are automatically hidden when they do not fit in the space between nodes to prevent overlapping of text and graphs; tree colours can be inverted for greater readability; node texts can be coloured according to node taxonomy.

Lastly, PhyD3 provides import and export tools to facilitate greater interoperability. Using the import tool users can supply trees in Newick and phyloXML formats, with optional numerical data, which can be easily converted to extended phyloXML format with graph annotations. Export capabilities support converting current tree visualisation into vector graphics (SVG format) and bitmaps (PNG format). A comparative overview of the main features of most popular phylogenetic tree viewers is given in Supplementary Table 1. A full feature list is described in Supplementary Table 2.

## 4. Conclusions and further information

We have developed an extension of the phyloXML standard to include graph specifications and datasets as extensions of the XSD scheme, as well as a phylogenetic tree viewer PhyD3, using the JavaScript D3.js library. Thanks to the phyloXML extensions, the PhyD3 viewer efficiently visualises extensively annotated phylogenetic tree data using modern web browsers. The software, full documentation, and demo material are available at http://phyd3.bits.vib.be and http://github.com/vibbits/phyd3

---

**Figure 1:**
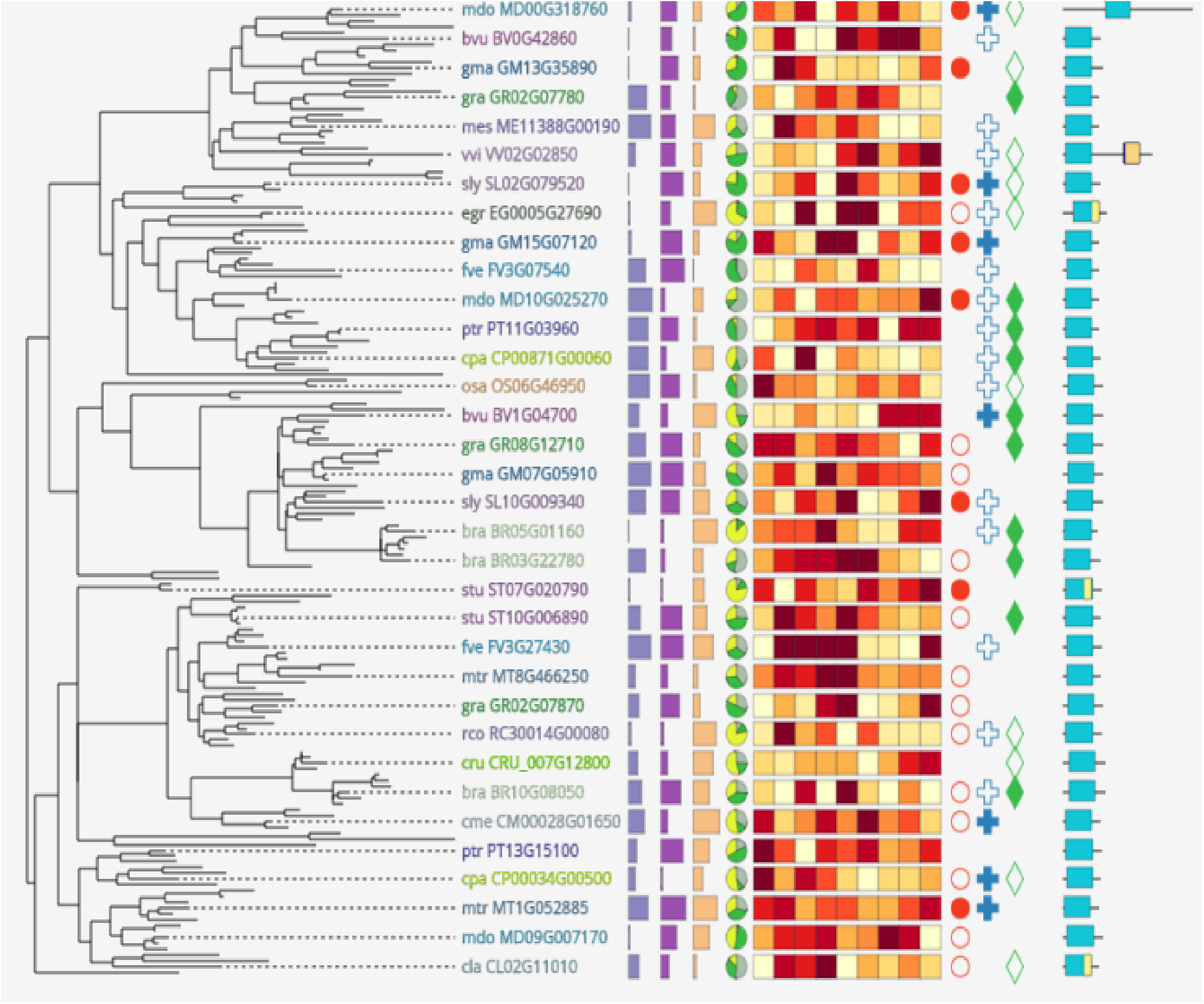
Example visualisation of phylogenetic tree using PhyD3

